# Lists with and without syntax: A new approach to measuring the neural processing of syntax

**DOI:** 10.1101/2020.05.18.101469

**Authors:** Ryan Law, Liina Pylkkänen

## Abstract

In the neurobiology of language, a fundamental challenge is deconfounding syntax from semantics. Changes in syntactic structure usually correlate with changes in meaning. We approached this challenge from a new angle. We deployed word lists, which are usually the unstructured control in studies of syntax, as both the test and the control stimulus. Three-noun lists (*lamps, dolls, guitars*) were embedded in sentences (*The eccentric man hoarded lamps, dolls, guitars*…) and in longer lists (*forks, pen, toilet, rodeo, graves, drums, mulch, lamps, dolls, guitars*…). This allowed us to perfectly control both lexical characteristics and local combinatorics: the same words occurred in both conditions and in neither case did the list items locally compose into phrases (e.g. ‘*lamps*’ and ‘*dolls*’ do not form a phrase). But in one case, the list partakes in a syntactic tree, while in the other, it does not. Being embedded inside a syntactic tree increased source-localized MEG activity at ~250-300ms from word onset in the left inferior frontal cortex, at ~300-350ms in the left anterior temporal lobe and, most reliably, at ~330-400ms in left posterior temporal cortex. In contrast, effects of semantic association strength, which we also varied, localized in left temporo-parietal cortex, with high associations increasing activity at around 400ms. This dissociation offers a novel characterization of the structure vs. meaning contrast in the brain: The fronto-temporal network that is familiar from studies of sentence processing can be driven by the sheer presence of global sentence structure, while associative semantics has a more posterior neural signature.

**SIGNIFICANCE STATEMENT:** Human languages all have a syntax, which both enables the infinitude of linguistic creativity and determines what is grammatical in a language. The neurobiology of syntactic processing has, however, been challenging to characterize despite decades of study. One reason is pure manipulations of syntax are difficult to design. The approach here offers a perfect control of two variables that are notoriously hard to keep constant when syntax is manipulated: word meaning and phrasal combinatorics. The same noun lists occurred inside longer lists and sentences, while semantic associations also varied. Our MEG results show that classic fronto-temporal language regions can be driven by sentence structure even when local semantic contributions are absent. In contrast, the left temporo-parietal junction tracks associative relationships.

## INTRODUCTION

Syntax is a combinatorial system that hierarchically relates linguistic elements during the construction of complex meaning. Its underlying neurobiology has been studied extensively for decades, yet a lack of consensus persists. One likely reason is a principled methodological challenge: it is very difficult to vary the syntactic structure of an expression without also altering its compositional semantics. Consequently, the nature and even existence of purely structural processing in the brain remains elusive. Here, we introduce a new experimental manipulation that succeeds in controlling certain semantic variables, specifically, word meaning and local semantic composition, better than prior research. With these robust modulators of neural activity controlled, will correlates of purely structural processing emerge?

Our study took advantage of the fact that word lists–typically used as unstructured control stimuli in studies of syntax–can also naturally occur inside a sentence, participating in the syntax of the sentence. For example, identical noun sequences can occur in longer lists and in sentences (Figure 1). In this contrast, the embedded three-noun lists across the pair of list-in-list and list-in-sentence are matched in their lexical characteristics and local combinatorics: in neither case do these words semantically or syntactically compose with one another (e.g. ‘*lamps*’ and ‘*dolls*’ do not form a phrase). A schematic depiction of this contrast is shown in Figure 1. In the list-in-sentence situation, the “syntactic engine” operates through the list, while in the list-in-list situation, it does not. This is our core contrast.

**Figure 1.**
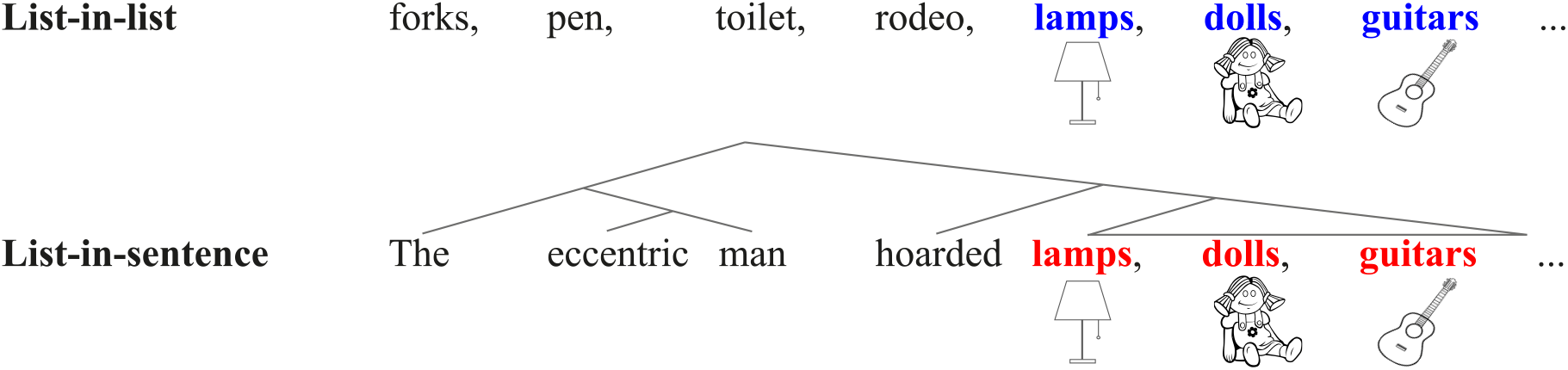
A schematic depiction of our structure contrast. The same noun list is embedded in a longer list (list-in-list) and in a sentence (list-in-sentence).

Since comparing lists and sentences—albeit not in the controlled fashion that we do here—has a long history in the neurobiology of sentence processing, that prior literature forms an important background for the current study. Emerging from this literature is a left lateral ‘combinatory network’ (Pylkkänen & Brennan, 2019) that is thought to subserve the composition of word meanings into larger syntactic and semantic structures: the anterior temporal lobe (ATL), the posterior temporal lobe (PTL), the inferior frontal cortex (IFC), the temporo-parietal junction (TPJ), and the orbitofrontal cortex (ORB). Summarised in Table 1, the findings point towards the left ATL as the most consistent correlate for sentence structure despite the variability in imaging techniques, stimulus presentation modalities, and types of unstructured controls. Targeted research on the ATL using magnetoencephalography (MEG) has, however, shown that the ATL does not appear to track syntactic structure building, but rather, aspects of conceptual combinatorics (Pylkkänen, 2019).

**Table 1.** A summary table showing a number of studies contrasting sentences to word lists using different imaging modalities, stimulus modalities, and control type.

| Studies | Imaging modality | Stimulus modality | Word list type | Language | ATL | PTL | IFC | TPJ | ORB |
| --- | --- | --- | --- | --- | --- | --- | --- | --- | --- |
| Mazoyer et al., 1993 | PET | Auditory | Content & function | French | × |  |  |  |  |
| Stowe et al., 1998 | PET | Visual | Content & function | Dutch | × |  |  |  |  |
| Vandenberghe et al. 2002 | PET | Visual | Sentence scrambled | English | × | × |  |  |  |
| Humphries, Love, Swinney, & Hickock, 2005 | fMRI | Auditory | Content only | English | × | × |  |  |  |
| Humphries, Binder, Medler, & Liebenthal, 2006; | fMRI | Auditory | Sentence scrambled | English | × |  |  | × |  |
| Snijders et al., 2009 | fMRI | Visual | Content & function | German | × | × | × |  |  |
| Pallier, Devauchelle, & Dehaene, 2011 | fMRI | Visual | Content & function | French | × | × | × | × |  |
| Brennan & Pykkänen, 2012 | MEG | Visual | Content & function | English | × | × | × |  | × |
| Fedorenko, Nieto-Castañón, & Kanwisher, 2012 | fMRI | Visual | Content & function | English | × | × | × | × |  |
| Matchin, Hammerly, & Lau, 2017 | fMRI | Visual | Content & function | English | × | × | × |  |  |
| Zaccarella, Schell, & Friederici, 2017 | fMRI | Visual | Content only | German |  | × | × |  |  |

The current study also used MEG, allowing us to measure reflections of structure position by position within our lists and to better understand the temporal nature of the effect. In addition, we also included a manipulation along a semantic dimension: association strength between the lists members. One possibility is that, regardless of the linguistic context in which the noun lists are embedded, the brain might (i) compose meaning when co-occurrence statistics between items are sufficiently high (Mollica et al., 2020), and/or (ii) more generally ‘chunk’ together nouns into some abstract representations (e.g. Christiansen & Chater, 2016) as a processing strategy (for instance, creating a coherent “grocery bag” scene from *juice, tomatoes, pasta*). Thus, by varying association strength, we aimed to distinguish potential effects of structure from effects of associative semantics. Crucially, our manipulation of association is distinct from manipulating semantic composition; to reiterate, members of noun lists do not combine with one another, neither syntactically nor semantically.

Neural responses were recorded as participants read word-by-word the same noun lists embedded in longer lists (unstructured controls) and in sentences (structured stimuli) then responded to a memory probe. Behaviorally, we would expect that the presence of structure facilitates recall (e.g. Potter et al., 1980, 2008). Neurally, we would expect the presence of structure to elevate cortical activation independent of word meaning if syntactic structure and lexical meaning can be dissociated; in contrast, we would expect comparable activity for lists in both structured and unstructured conditions if syntax and word semantics cannot be dissociated.

## MATERIALS AND METHODS

### Stimuli & Design

We selected concrete English nouns based on the concreteness rating corpus by Brysbaert, Warriner, and Kuperman (2014). From this pool, we then selected nouns that are matched in their log frequency from the SUBTLEX-US corpus (Brysbaert & New, 2009). The critical list nouns were changed from their singular form to plural to block potential noun-noun compounding (e.g. *lamp doll* could form a phrase, but *lamps dolls* could not). These plural nouns were then used to construct our critical three-noun lists such as *lamps, dolls, guitars*. The lexical characteristics are summarized in Table 2.

**Table 2.** Lexical characteristics of the critical items. Mean values for each measure are reported, with SD in parentheses.

| Condition | Log Frequency | Concreteness | Word length | Sentence Plausibility | Association Strength |
| --- | --- | --- | --- | --- | --- |
| **List-in-list, Low Association** | 2.45 (0.65) | 4.87 (0.13) | 6.02 (1.68) | - | 0.44 (0.09) |
| **List-in-sent, Low Association** |  |  |  | 5.95 (1.40) | 0.45 (0.09) |
| **List-in-list, High Association** | 2.59 (0.63) | 4.84 (0.18) | 5.56 (1.49) | - | 0.59 (0.10) |
| **List-in-sent, High Association** |  |  |  | 6.18 (1.30) | 0.62 (0.09) |

For lists-in-lists, the nouns surrounding the critical lists were assigned at random. We prepended four and appended three nouns to the critical lists, resulting in ten-word sequences. The number of plural, singular mass, and singular count nouns was balanced. Items from the critical lists were included in other items as non-critical nouns (i.e. filler nouns surrounding critical lists) to balance out the co-occurrence statistics of the critical items within our stimulus set. As for lists-in-sentences, the same critical lists were given a sentence frame. We prepended a subject and a verb, as well as appended two additional nouns connected with the conjunction *and* to the end of critical lists, resulting in ten-word sentences. The presence of a determiner preceding the critical lists was balanced across conditions.

For the association strength manipulation, we calculated word co-occurrence statistics by first extracting vectors of the stimuli content words from a pre-trained Global Vectors model (Pennington et al., 2014). Then, we calculated the cosine similarity of content words across Words 1 to 7 and made sure that the distribution of association strength was bimodal, with high and low association cases reflecting each of the local maxima. The sentences were then submitted to a norming survey for plausibility on Amazon Mechanical Turk. A stimulus set thus consisted of four ten-word sequences: two lists-in-lists and two lists-in-sentences. Set-hood was defined here by a common word 7 (e.g. *guitars*) instead of words 5 and 6 so as to allow us to vary those words and subsequently varying association strength. An example stimuli set is shown in Table 3. In total, the experiment comprised of 168 trials. All stimuli are shown in the appendix.

**Table 3.** One complete set of stimuli showing the full 2×2 design crossing Structure and Association.

| Structure | Association | Words 1-4 | Words 5-7 | Words 8-10 |
| --- | --- | --- | --- | --- |
| **List-in-list** | **Low** | forks pen toilet rodeo | **lamps dolls guitars** | wood symbols straps |
| **List-in-sent** | **Low** | The eccentric man hoarded | **lamps dolls guitars** | watches and shoes. |
| **List-in-list** | **High** | theatre graves drums mulch | **pianos violins guitars** | crates knuckle cocoa |
| **List-in-sent** | **High** | The music store sells | **pianos violins guitars** | drums and clarinets. |

### Experimental procedures

Stimuli were delivered using rapid serial visual presentation of white text on grey background back-projected onto a monitor about 80cm away from participants’ head. Participants initiated each trial via a button press. Each trial began with a fixation cross on screen for 300ms, followed by an inter-stimulus interval (ISI) of 300ms. Stimulus words were also presented on screen for 300ms then an ISI for 300ms. At the end of each trial, a memory probe appeared on screen, consisting of a word in blue. Participants responded to the task via button press: they pressed the left button if that word was drawn from that trial, the right button if not (Figure 2). Behavioral reaction times and accuracy scores were measured from the presentation of the memory probe task. Items were fully randomized across the experiment.

**Figure 2.**
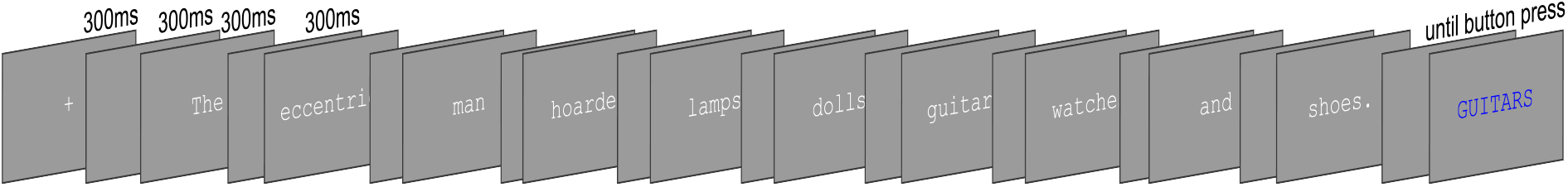
Trial structure.

The memory probe task was selected for both lists-in-lists and lists-in-sentences to monitor participants’ attention. A random word (either content or function) was drawn pseudo-randomly from each trial. Although the task was likely less demanding for lists-in-sentences, as sentences might be privileged in working memory (Baddeley et al., 2009) and therefore were expected to be easier than lists-in-lists (Potter et al., 1980, 2008), adopting a parallel task across conditions was deemed more important. In fact, under the assumption that harder processing engages the brain to a greater extent, having a word recall task might increase the brain’s responses to the unstructured list-in-list conditions. Furthermore, as aforementioned, a possibility remains that participants work to create phrases out of word lists or use chunking as a strategy when faced with a difficult task. Together, the experimental conditions might reduce the difference in cognitive operations engaged by the two conditions and thus biasing our study against finding an activity increase for lists-in-sentences over lists-in-lists, which was the effect of interest.

### Participants

Twenty-two native English speakers participated in the experiment. Two were excluded due to technical issues during data acquisition; four were excluded due to excessive sensor noise. Thus, a total of sixteen participants were included in our analyses (9 women; mean = 24.8 years, SD = 7.4 years). All participants are right-handed and reported no history of neurological disorder.

### Data acquisition and pre-processing

Before recording, each participant’s head shape was digitized using a Polhemus FastSCAN system (Polhemus, Vermont, USA). Digital fiducial points were recorded individually, including three anatomical landmarks (the nasion and the left and right tragi) and five marker coil positions (three points on the forehead and one point each at 1cm anterior to the left and right tragi). Marker coils were placed at the same five positions in order to localize the participant’s head relative to the MEG sensors. The measurements of head position using marker coils were recorded right before and after experiment to correct for movement during recording. MEG recordings were collected in the MEG Lab at New York University Abu Dhabi using a whole-head 208 channel axial gradiometer system (Kanazawa Institute of Technology, Kanazawa, Japan) as participants lay supine in a dimly lit, magnetically shielded room. A practice session first took place outside the magnetically shielded room.

MEG recordings were sampled at 1000Hz with an online band-pass filter between 0.1Hz-200Hz and noise reduced using eight reference channels via the Continuously Adjusted Least-Squares Method (Adachi, Shimogawara, Higuchi, Haruta, & Ochiai, 2001), in the MEG Laboratory software (Yokogawa Electric Corporation and Eagle Technology Corporation, Tokyo, Japan). The noise-reduced MEG recording, the digitized head-shape and the head position measurements were then imported into MNE-Python (Gramfort et al., 2014). Data were submitted to an offline low-pass filter of 40Hz with a finite impulse response filter design using a Hamming window method. Flat or excessively noisy channels were interpolated using the spherical spline method (Perri, Pernier, Bertrand, & Echallier, 1989). The data was then submitted to an independent-component analysis for detection and removal of well-characterized artefacts (eye blinks and heart beats) and noise components characteristic of the MEG system. Finally, data were segmented into epochs spanning the whole ten-word sequences, each baselined using the 200ms period prior to trial onset. Epochs were automatically rejected if any sensor value after noise reduction exceeded 2.5 pT/cm at any time. Then, epochs were trimmed to contain only the critical list items.

We estimated cortical activity by creating dynamic statistical parameter maps (dSPM: Dale et al., 2000). First, MEG data were co-registered with either the participant’s anatomical MRI when available or the FreeSurfer average brain when not (CorTechs Labs Inc., California, USA and MGH/HMS/MIT Athinoula A. Martinos Center for Biomedical Imaging, Massachusetts, USA). The FreeSurfer average brain was scaled to match the participant’s head-shape while aligning the fiducial points. Minute manual adjustments were conducted to minimize the difference between the head shape and the average brain. Next, a source space was set up, each hemisphere containing 2562 potential electrical sources. A forward solution was then computed using the boundary element model. Channel noise covariance matrices were estimated using the baseline period (200 ms before trial started) and regularized using the automated method (Engemann & Gramfort, 2015). Combining the forward solution and noise covariance matrices, an inverse solution was computed and applied to participant evoked responses assuming a free orientation of the current dipole to yield cortical source activity estimates.

Our primary analyses were performed on source activity localized to the five regions of interest (ROIs) from each hemisphere (Figure 3). The inclusion of right hemisphere homologues was motivated by findings showing right hemisphere involvement during combinatory language understanding (Mazoyer et al., 1993; Stowe et al., 1998; Humphries et al., 2006; Rogalsky & Hickok, 2009). The ROI labels are defined and generated as follows. For the left IFC, ATL and PTL, 30 mm spheres were created around coordinates for the left inferior frontal cortex, temporal pole, anterior superior temporal sulcus, posterior superior temporal sulcus in Montreal Neurological Institute (MNI) space reported by Pallier et al. (2011). Spheres for the left temporal pole and anterior superior temporal sulcus were combined, because studies often find these regions co-activating, and because MEG is less spatially resolved compared to fMRI. To generate the right hemisphere homologues of these regions, the polarity of the x-axis values in the MNI coordinates were flipped before 30 mm spheres were created around them. Both left and right TPJs were generated by combining the angular gyrus (Brodmann area 39) and adjacent supramarginal gyrus (Brodmann area 40) labels from the “PALS-B12-Brodmann” atlas (Van Essen, 2005), while left and right ORBs were generated by combining lateral and medial orbitofrontal labels from the “aparc” atlas (Desikan et al., 2006).

**Figure 3.**
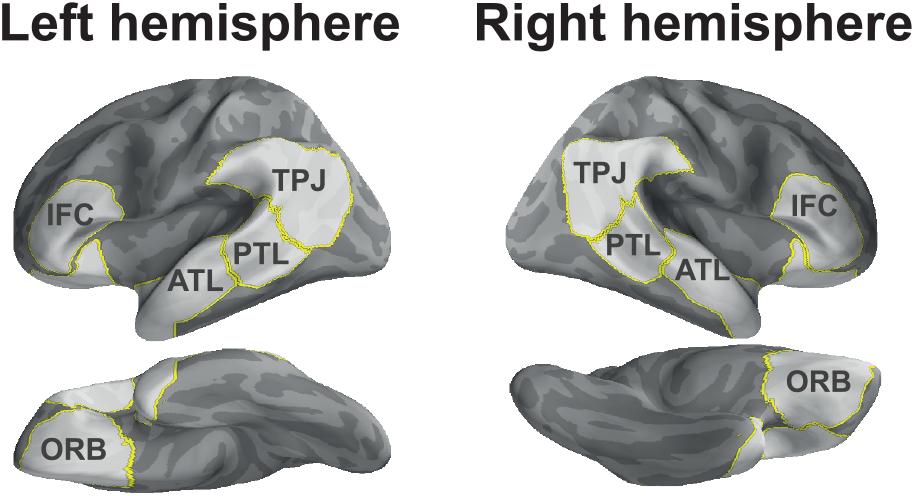
Regions of interest

### Statistical Analysis

#### ROI analysis

We performed temporal and spatiotemporal non-parametric cluster-based permutation tests (Maris & Oostenveld, 2007) using the python package Eelbrain (v0.30.11; Brodbeck, 2019) in each ROI and across the whole brain respectively (see Bemis & Pylkkänen, 2011 for a similar application). For temporal tests, source-localized MEG estimates were first averaged across sources within each ROI. Since all critical list items participate in their larger surrounding context in the same way, we included word position (words 5, 6, and 7) as a factor. This allowed us to examine potential structure effects time-locked to word presentation and afforded us better statistical power. Then, a 2×2×3 repeated measures analysis of variance (ANOVA) was fitted at each time sample separately across the whole epoch (600 ms). Factors included structure (list-in-list, list-in-sentence), association (high, low), as well as position (word5, word6, word7). Temporal analyses adopted a cluster forming threshold of *p*<.05 with a minimum of 20 contiguous time samples. For the spatiotemporal test, we followed the same procedure, but instead performing a spatiotemporal search across the whole brain, i.e. without averaging activity across source space; In this analysis, an additional criterion of cluster forming threshold with at least 20 contiguous spatial samples was adopted. Cluster level *p*-values were first estimated via Monte Carlo simulations, repeated 10,000 times. These *p*-values were then corrected for multiple comparisons across all ROIs by controlling the false discovery rate at the critical value of 0.05 (FDR; Benjamini & Hochberg, 1995; Genovese, Lazar, & Nichols, 2002)

## RESULTS

### Behavioral results

For each participant, we removed reaction time measures that either corresponded to incorrect responses or were two standard deviations from their own mean. Next, we fitted a linear mixed effects regression model to log-transformed reaction time data using the *lme4* package in R (Bates et al., 2015). The model included fixed effects for structure (list-in-list, list-in-sent) and association (continuous association measures). The model also included a full random effects structure over subjects for all fixed effects and random intercepts for items. We found a main effect of structure (*χ*^2^=44.73, *p*<.001). Compared with the corresponding list-in-list conditions, participants were markedly faster for lists-in-sentences (mean ± SD, 965 ± 294 ms) relative to lists-in-lists (828 ± 249ms). Association strength between list items and contexts did not modulate reaction times (*p*=.877). We then fitted a generalized mixed effects logistic regression to the accuracy data (incorrect responses included). Again, we observed a main effect of structure (*χ*^2^ =20.24, *p*<.001) but not association (*p*=.425). Participants’ responses were more accurate to lists-in-sentences (96% ± 18%) compared to lists-in-lists (88% ± 31%).

As expected, we found that structure facilitated recall. Here, we explore two possible explanations. First, sentences are much more engaging as stimuli than long lists of nouns in general. Thus, participants might have paid more attention to sentence stimuli, leading to better performance as a result. Second, although both structure types contain the same number of words, the likelihood of drawing a noun is higher in lists-in-lists (which consisted only of nouns) compared to lists-in-sentences. As such, there were more noun competitors for lists-in-lists in the word recall task, which may subsequently reduce response accuracy and increase reaction times. Importantly though, the present behavioral results demonstrate that the word recall task was *less* demanding for conditions *with* sentence structure. Under the assumption that an easier task engages the brain to a lesser extent, any increase in our structured stimuli could not be attributed to general increased task effort.

### Neural effects of structure

In the left PTL, a cluster extended from 333 to 389 ms after word onset. The cluster-based permutation test indicated that there was a significant effect of structure (*p*=.004). Inspecting activity waveforms, lists in a structured expression increased PTL activity relative to lists in a longer list (Figure 4). Another cluster spanned from 66 to 93 ms in this region. The permutation test implicated a structure by position interaction (*p=*.037). Pairwise comparisons revealed that word 5 from lists-in-lists and word 7 from lists-in-sentences elicited significantly stronger activity than word 7 in lists-in-lists. Turning to the left IFC, the structure effect was driven by a sharp increase in cortical signals elicited by structured lists relative to unstructured ones, as revealed by the permutation test in this region (*p*=0.028). This effect corresponded to a cluster spanning 237-272 ms post word onset. Within the left ATL ROI, the permutation test revealed significant structure effects (*p*=.003, *p*=.024), which corresponded to two clusters. A cluster emerged very early at 0-51 ms post word onset, while a second cluster at 312-357 ms. In both of these clusters, lists-in-sentences elicited greater activity than lists-in-lists. A closer inspection at activity time-locked to each list item suggested that ATL activity was elevated for structure in a sustained fashion. As a result, that early waveform separation might be a result of averaging sustained activity across the lists. The main effects of structure were broken down by the full design in Figure 5.

**Figure 4.**
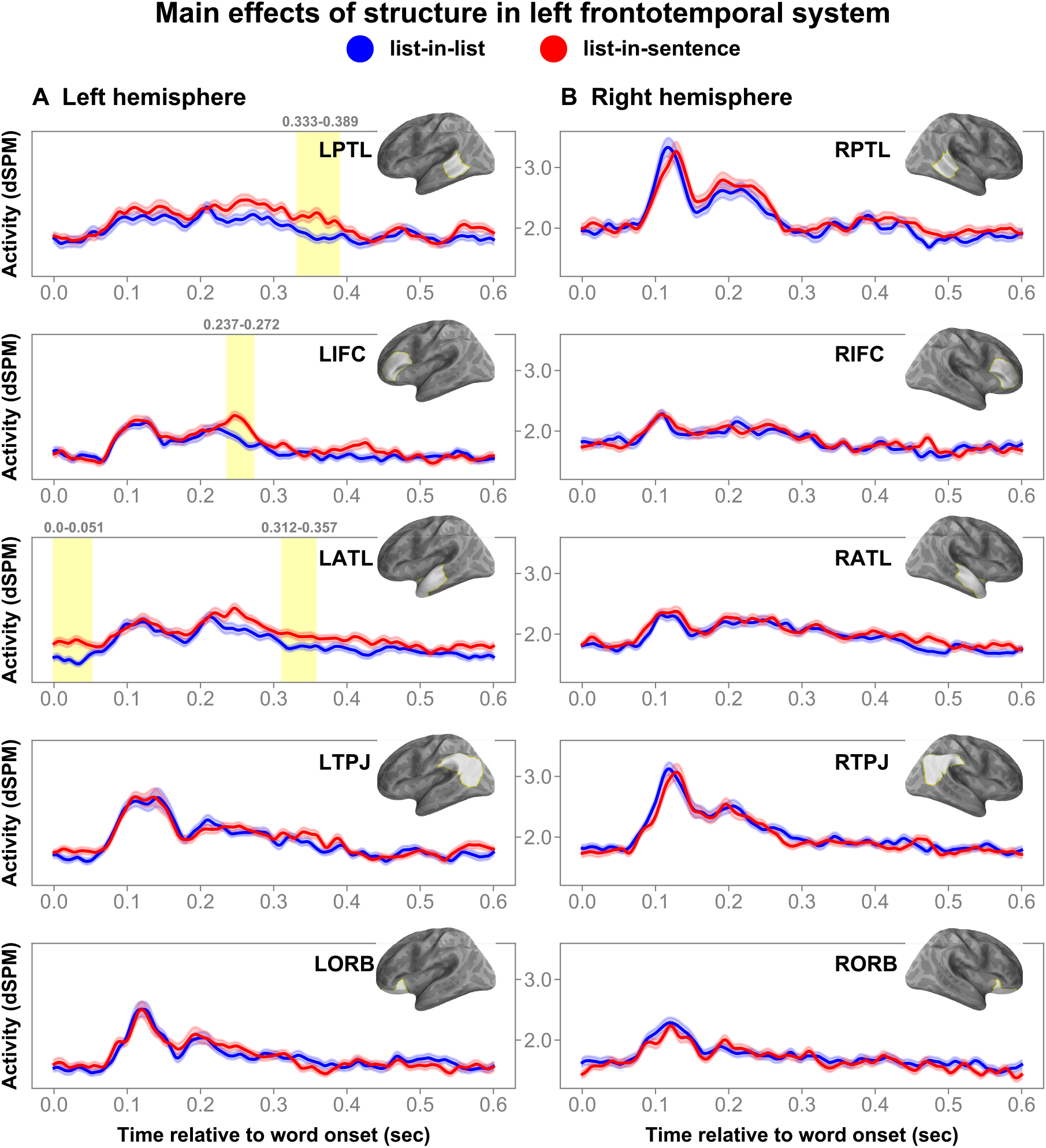
Main effects of structure. Panels (A) and (B) chart the time courses in the left and right hemispheres respectively. The brain model indicates the ROI analyzed. Shaded regions in timeseries indicate cluster extent corresponding to FDR corrected significant effects at p-values less than 0.05. Error bars represent one within-subjects standard error from mean (Loftus & Masson, 1994).

**Figure 5.**
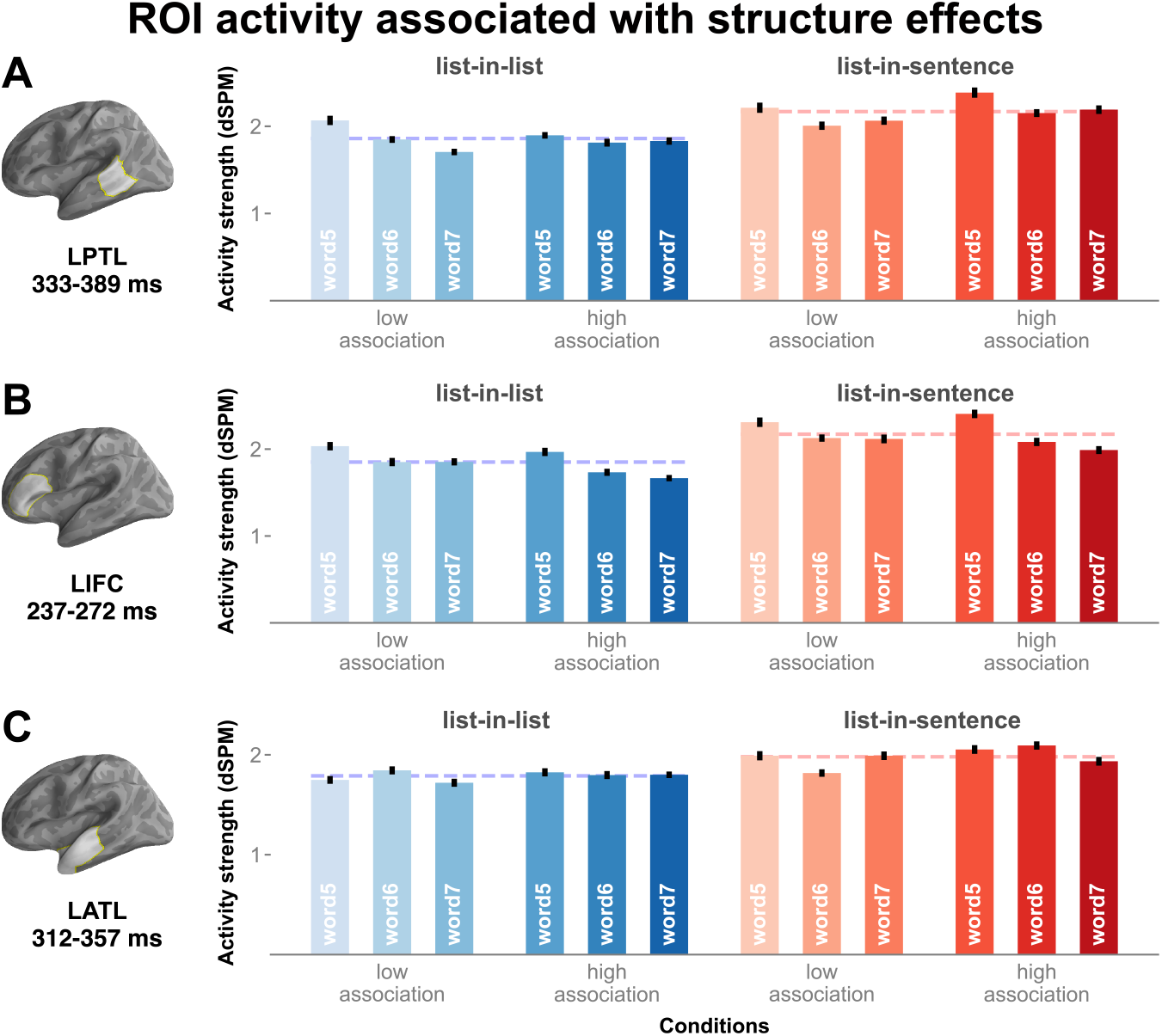
Panels (A), (B), and (C) show averaged cluster activity by the full design in the left PTL, left IFC, and left ATL respectively. Activity elicited by lists-in-lists are colored in blue, while lists-in-sentences red. Dashed lines represent averaged cluster activity elicited by lists-in-lists and lists-in-sentences. Error bars represent one within-subjects standard error away from mean (Loftus & Masson, 1994).

In the *right* ORB, although there was no main effect of structure, structure and association did interact (*p*=.027). This interaction was captured in a cluster at 176-199 ms post word onset. ROI activity was higher for lists-in-lists with high association than low association; this contrast in activity was not significant within lists-in-sentences.

### Structure by region interaction

To directly assess whether we can reject the null hypothesis that there is no significant difference between structured vs. unstructured cases in other regions within the left hemisphere, source activity from left hemisphere ROIs were submitted to an omnibus 2⨉2⨉3⨉5 (structure by association by position by region) ANOVA, with the same test criteria outlined above, across the left hemisphere. The analysis indicated a significant structure by region interaction (*p*=.008). Post-hoc pairwise comparisons of averaged activity within this time window suggest that the observed difference between lists-in-lists and lists-in-sentences was largest in the left PTL ROI, with lists-in-sentences driving higher activity in this ROI (Figure 6A). The corresponding cluster extended from around 330 to 380 ms, which matched the cluster observed in the left PTL in the ROI analysis described above. Though there were significant structure effects in the left IFC and left ATL, their respective clusters appeared to have different temporal profiles than that observed in the left PTL.

**Figure 6.**
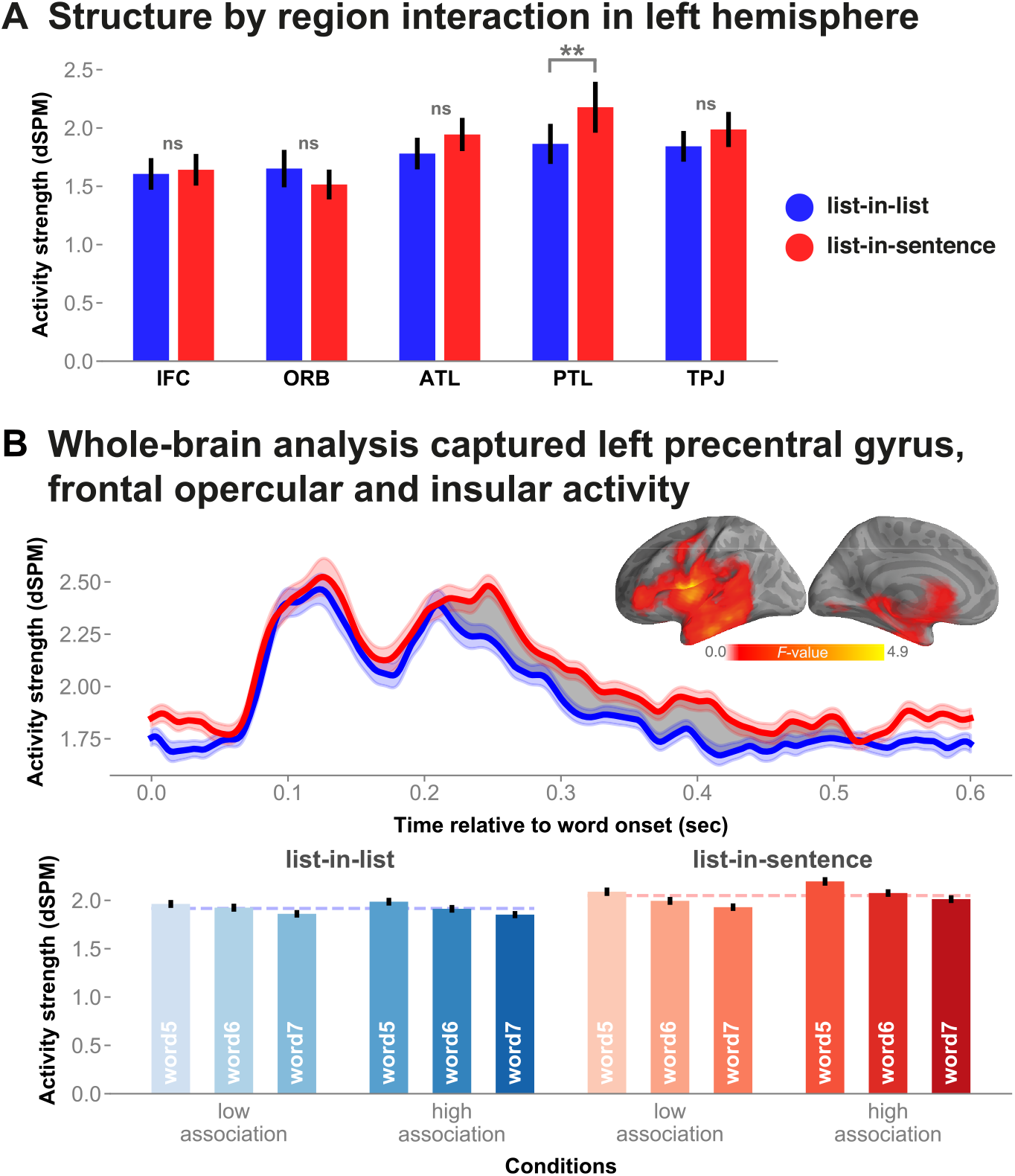
Panel (A): Structure by region interaction. Bar plot shows averaged cluster activity at 332-381 ms post word onset in all left hemisphere regions. Panel (B): shows the main structure effect indicated in the spatiotemporal cluster-based permutation analysis across the whole brain. The shaded area in the time series marks the cluster’s temporal extent, while the brain model shows the spatial extent. The red-blue bar graphs break down the main structure effect by the full design. ns, Non-significant, **p<.01)

In addition, we ran a complementary cluster-based spatiotemporal analysis across the whole brain. The analysis indicated a significant effect of structure (*p*=.0017): increased activity for structured lists relative to unstructured controls. A cluster extended temporally from 189 to 508 ms post noun onset. (Figure 6B). Spatially, the cluster extent covers the left frontotemporal regions, portions of the left frontal operculum, the underlying left insula, as well as portions of the left precentral gyrus.

### Neural effects of association

A cluster extended from 375 to 413 ms after stimulus onset in the left TPJ. The permutation test in this region indicated a main effect of association (p=.031) (Figure 7). Lists that were more strongly associated in terms of co-occurrence elicited stronger signals than lists in contexts that are relatively less associative. Examining the activity waveform, the cluster extent encompassed a peak in amplitude at around 400 ms post word onset, a timing that bore semblance to that of the N400 event-related potential component. Averaged cluster activity time locked to each list item was plotted in bars in (Figure 7B). There was no significant association by position interaction.

**Figure 7.**
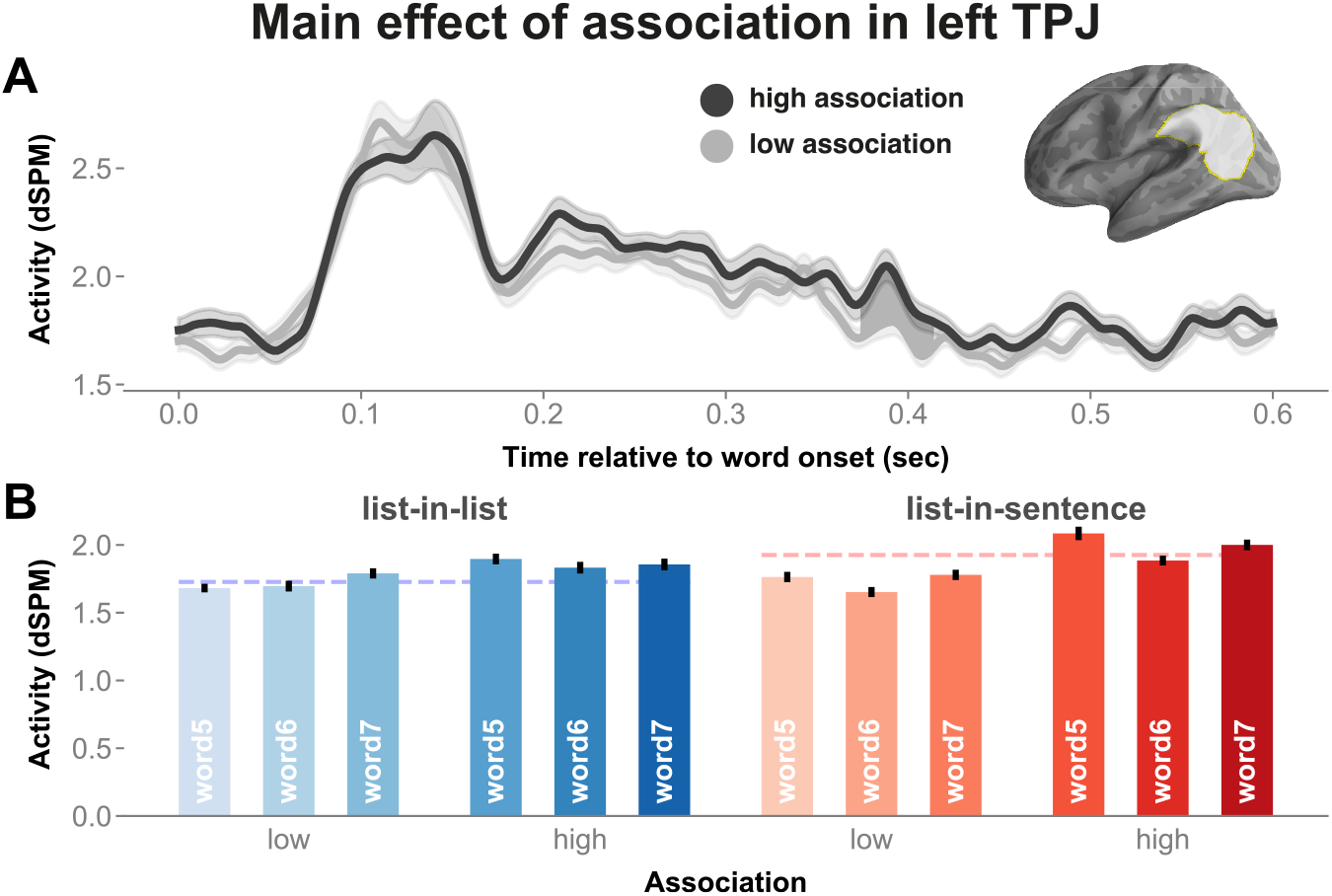
Panel (A) charts the activity time course in the left TPJ, with cluster extent shaded in grey. Panel (B) shows the red-blue bar graphs, breaking down the effect of association by the full design. Error bars represent one within-subjects standard error from mean (Loftus & Mason, 1994).

### Neural effects of position

Across the ROI and spatiotemporal analyses, we also observed position effects (Figure 8). While this is not the main effect we sought to interpret, it is interesting to note that activity was greater for Word 5 than Words 6 and 7 across lists with and without structure. This pattern of activity was observed across a broad set of regions in both hemispheres.

**Figure 8.**
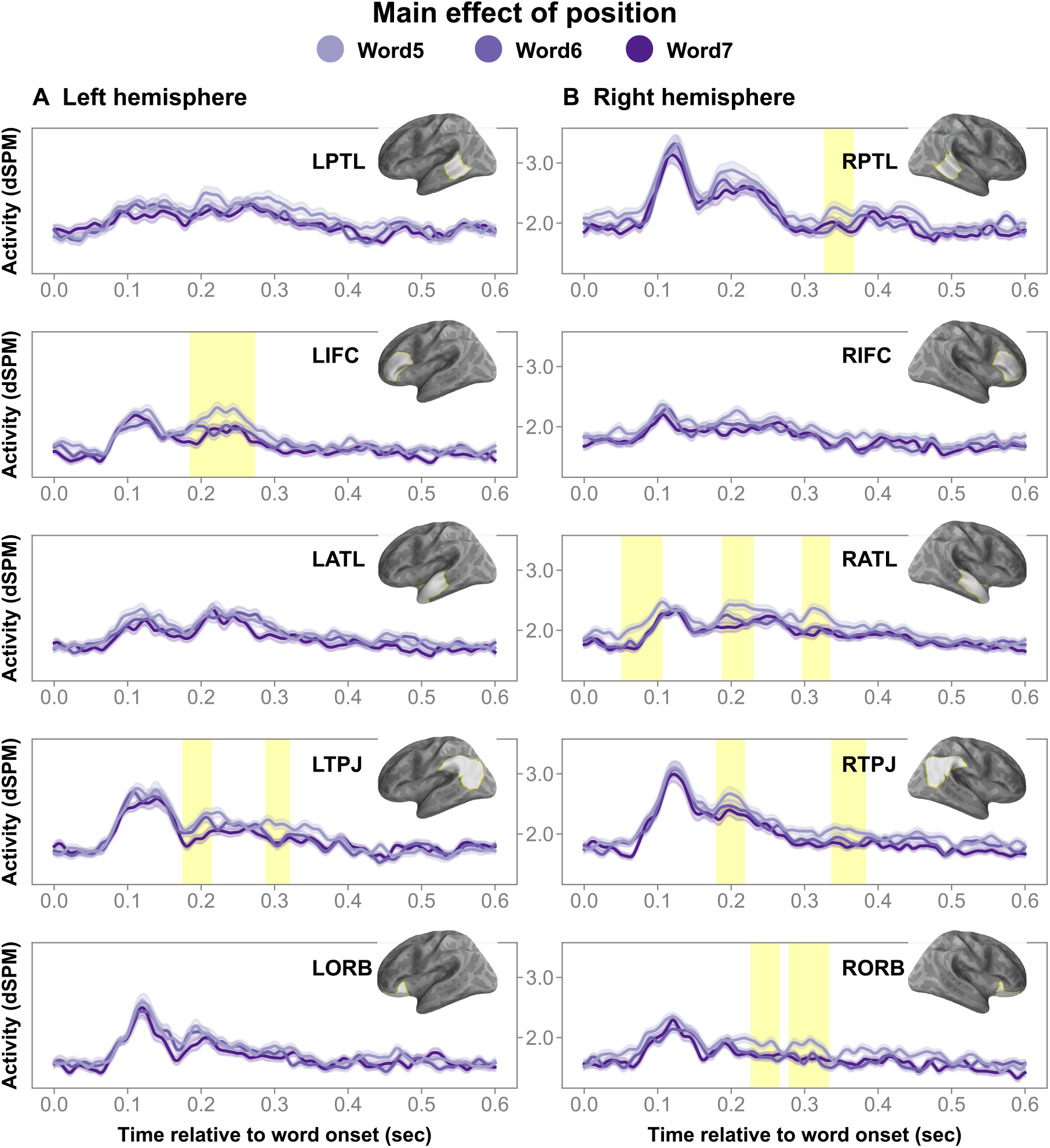
Main effects of position. Panels (A) and (B) chart the time courses in the left and right hemispheres respectively. The brain model indicates the ROI analyzed. Shaded regions in time series indicate cluster extent corresponding to FDR corrected significant position effects.

Indeed, within a list embedded in a structured expression, word 5 (i.e. the first member of the list) is the first noun that the verb takes as a direct object (e.g. *The eccentric man hoarded lamps…*). This invites the question of whether the increase in activity we observed for lists-in-sentences were due to processes associated with the integration of the direct object with the verb (although obviously within our stimuli, the argument slot is not fully saturated until the rest of the conjunction is processed). Therefore, a possibility remains that the structure effects we observed above were largely driven by argument structure related processes associated predominantly with word 5.

To investigate this possibility, we excluded word 5 and performed cluster-based permutation tests across all ROIs (and corrected for multiple comparisons accordingly) with the same test criteria outlined above. In other words, we collapsed across words 6 and 7, resulting in a 2⨉2⨉2 ANOVA at each time sample in each ROI. We found that the structure effects held in the left PTL (*p*=.01) and left ATL (*p*<.01). The IFC’s effect of structure now only approached significance (*p*=.08, uncorrected). The corresponding clusters’ extents resembles their three-word counterparts in size and timing to a large extent. Thus, the reduction in IFC structure effect is likely a result of a reduction in statistical power from removing word 5.

## DISCUSSION

As a novel approach to the neurobiology of syntax, we devised an MEG experiment that identified neural correlates of structure while controlling many related, hard-to-control stimulus properties. Within the combinatory network, the left PTL, left IFC and left ATL demonstrated sensitivity to the presence of structure. These structure effects cannot be attributed to (i) contributions from word-level semantics (the same words occurred at the same time points within our structure contrast), (ii) local phrase combinatorics (the critical list items were all plural, blocking noun-noun compounding), (iii) general task demands (at least under the usual assumption that more effort leads to higher neural activity—here, activity was higher in cases where recall was facilitated). Main effects of structure corresponded to clusters emerging at different time points, providing the temporal resolution lacking in previous hemodynamic work.

### Left posterior temporal lobe

Recent MRI and MEG findings suggest several hypotheses about the functional role of the left PTL in the context of combinatory processing. First, the PTL is thought to play a critical role in lexical storage and retrieval (for a review, see e.g. Lau et al., 2008). It has been suggested that lexical items are stored together with their associated semantic content as well as syntactic information (e.g. Snijders et al., 2009; Rodd et al., 2010; Tyler et al., 2013; Matchin et al., 2017; Matchin, Brodbeck, et al., 2019; Matchin, Liao, et al., 2019). One of our critical findings is that when the semantic content has been matched across levels of structure, we still observed effects of structure: PTL activity was higher for the same word sequences embedded in a structured context compared to the longer, unstructured counterpart. Furthermore, a structure by region interaction across the left hemisphere language-related areas suggested that the structure effect was the strongest in the left PTL at around 330-380 ms. While the PTL’s contribution in lexico-syntactic access is evident from the literature, the present study suggests that, at a minimum, partaking in a syntactic tree drove the left PTL above and beyond the process of accessing lexical representations and their associated syntactic information.

Second, a recent MEG study demonstrated the PTL’s early contribution in syntactic composition, as evidenced by contrasting two cases in which semantic composition took place in both cases but syntactic composition only in one (Flick & Pylkkänen, 2020). Our study also sought to isolate syntax but contrasts with Flick and Pylkkänen (2020) in that our critical list items did not compose with one another, thus minimizing cortical activity associated with local compositional semantics within our critical regions. Matching content and blocking conceptual combination, our study found increased PTL activation at around 330 to 390 ms, while Flick and Pylkkänen (2020) at around 200 to 230 ms.

The PTL was also recently proposed to engage in predicting likely upcoming hierarchical syntactic structures (dubbed the structural prediction account; see Matchin, Brodbeck, et al., 2019; Matchin et al., 2017). However, a possibility remains that semantic factors are likely also brought to bear on predictions online (see e.g. Kuperberg & Jaeger, 2016). Thus, more work is needed to rule out semantic contributions to structural prediction. Moreover, studies finding structural predictive effects (e.g. Matchin, Brodbeck, et al., 2019) adopted a blocked design, which would encourage prediction. Given that the stimuli in the present study were fully randomised, our results are possibly partly accounted for by a structural prediction account of the left PTL.

### Left inferior frontal cortex

Activity increase associated with the presence of structure was observed in the IFC at around 240 to 270 ms with a more distinctly evoked nature. This finding contrasts with PTL finding, wherein increased activity had a more sustained quality at around 330 to 390 ms. In fact, examining the interaction between structure and region suggests that within the PTL structure effect time window, IFC activity was comparable across structures (Figure 6). With caution, our interpretation of the pattern of results is that the left PTL and IFC likely carry out distinct functions with regards to syntactic processing, both regions working in tandem during combinatory language comprehension (e.g. Griffiths et al., 2012; Tyler & Marslen-Wilson, 2008).

### Left anterior temporal lobe

The composition of sentence meaning amounts to a cascade of processes at multiple levels of representation (syntactic, semantic, pragmatic, etc.). Thus, a body of work has sought to strip away sentence-level computations in order to unpack the constituent processes and investigate the neural reflexes of meaning composition in simple two-word phrases (for a review, see Pylkkänen, 2019). The composition of two-word phrases correlates with an increase in ATL activity around 200ms post noun onset. Although originally compatible with the hypothesis that this activity is reflective of syntactic structure building (Bemis & Pylkkänen, 2011), subsequent studies have refined our understanding of the ATL’s function. In particular, ATL activity during meaning composition was shown to be sensitive to semantic features, such as the conceptual specificity of the constituent items across expressions with comparable syntax (e.g. comparing pairs like [blue boat, blue canoe] or [meat dish, lamb dish]; see Westerlund & Pylkkänen, 2014; Zhang & Pylkkänen, 2015). Whereas, when comparing two expressions that have relatively parallel conceptual content but divergent syntactic combination, ATL remains insensitive (Flick & Pylkkänen, 2020). There are also preliminary results that the combinatorial steps underlying meaning composition in adjective-noun pairs are largely insensitive to syntactic structure (Kim & Pylkkänen, 2020; Parrish & Pylkkänen, 2019).

Within the critical lists, we controlled for the local composition of phrases (i.e. the list items do not compose with one another to form phrases). This lack of local phrasal composition would predict the absence of a combinatory left ATL effect. Contrary to this prediction, our left ATL results showed a main effect of structure. This finding suggest that the left ATL’s participation during combinatory sentence understanding goes beyond accessing individual word meanings and combining the denoted conceptual features. Our current results cast a new light on the left ATL’s function, opening up a new research question about this region. To the extent that combinatory syntax and semantics can be dissociated, a pertinent question, then, is whether the ATL activity observed in our study was reflective of sentence-level syntax or sentence-level semantic interpretation. In sum, the presence of a structure effect and the absence of an association effect within the left ATL rule out explanations in terms of lexical (semantic and/or syntactic) access and local semantic composition.

### Left temporo-parietal junction: Associative semantics

We found a main effect of association in the left TPJ: stronger association among list items and contexts elicited stronger activity. Although the TPJ effect is in the N400 time window (Figure 7), and it is an effect of a stimulus factor that classically affects the N400, the directionality of our effect is the opposite from what is expected for the N400. When words are primed with associated words, N400 amplitude is expected to *lower* (e.g. Ortu, Allan, & Donaldson, 2013; Rhodes & Donaldson, 2008). By contrast, in our results, source amplitude for words in more associative contexts *increased*. (see Kutas & Federmeier, 2011 for a general review of the N400).

Importantly, activity housed in the left TPJ increased as a function of semantic association strength regardless of structure. A tentative hypothesis might be that, as levels of association strength increased, the brain might attempt to ‘make sense’ out of the plural nouns. This computation might take the form of (a combination of) semantic composition (e.g. Mollica et al., 2020) and chunking (e.g. Christiansen & Chater, 2016). For example, the highly associative list “*pianos, violins, guitars*” might result in a grouping of a “musical instrument” set compared to a less associative list “*lamps, doll, guitars*”. In all, the left TPJ is the only region that showed sensitivity to our association contrast, and critically, no sensitivity to the structure contrast. This suggests a more conceptual-semantic function as opposed to a syntactic one for the TPJ, consistent with previous proposals (e.g. Matchin, Brodbeck, et al., 2019; Pallier et al., 2011).

### Limitations and future directions

Throughout this article, we have alluded to the fact that a confound exists in our design: lists-in-sentences introduce a global semantic context that describes a scenario, while lists-in-lists do not. However, our design succeeds in controlling semantics at a more local level. Individual word meanings and phrasal composition are perfectly matched in our paradigm – an approach never seen in prior research. Therefore, we believe the current paradigm, appropriately refined, would be suitable to future work for distinguishing the contributions of sentence-level message vs. sentence-level syntactic structure. Further, this paradigm can also be extended to other syntactic categories such as adverbs, adjectives, and verbs. The present study used sequences of nouns. It is possible that our set of results does not generalize beyond noun lists. Thus, a straightforward follow-up would be to extend this paradigm to such other syntactic categories and draw up lists accordingly (e.g. *the big blue wooden ball*).

## CONCLUSION

The present study leveraged word lists, which are typically used as unstructured controls, to deconfound syntax from semantics. We found that the left PTL, left IFC, and left ATL showed sensitivity to structure independent of lexico-conceptual semantics and local combinatorics. A structure by region interaction indicated that the observed difference between lists-in-lists and lists-in-sentences was largest in the left PTL. While explanations in terms of the global semantics of the sentences cannot yet be ruled out, this pattern of results allowed us to rule out explanations in terms of lexical semantics and local semantic composition. Our left TPJ finding supports a conceptual-semantic role for the region. These findings contribute a piece to our relatively coarse understanding of the extent to which syntactic and semantic processes could be teased apart at the level of brain regions, as well as providing the temporal resolution that hemodynamic work lacks.

## Acknowledgements

This work was funded by the New York University Abu Dhabi Institute Grant G1001 (L.P.).

## Appendix test stimuli

Appendix: test stimuli
| Set | Structure | Association | Item |
| --- | --- | --- | --- |
| 1 | sent | low | Tony just bought some pasta, soap, milk, salt, and soda. |
| 1 | list | low | yard, cello, desk, stadium, pasta, soap, milk, charcoal, stalls, pebbles |
| 1 | sent | high | The tasty dish contains butter, cheese, milk, garlic, and spinach. |
| 1 | list | high | bell, dryer, goats, tuna, butter, cheese, milk, nuts, salt, sugar |
| 2 | sent | low | Luckily Phil brought some tissues, raisins, cups, plates, and chips. |
| 2 | list | low | cart, corridors, yachts, chives, tissues, raisins, cups, feathers, fan, freezers |
| 2 | sent | high | Finally Sam remembered the bowls, spoons, cups, sandwiches, and forks. |
| 2 | list | high | shelf, sushi, polyester, coin, bowls, spoons, cups, windows, cookies, seals |
| 3 | sent | low | The helpers prepared some broth, fish, bread, crabs, and batter. |
| 3 | list | low | cabinets, mirrors, thigh, pool, broth, fish, bread, honey, deserts, desks |
| 3 | sent | high | The waiter served some pizza, salad, bread, butter, and soup. |
| 3 | list | high | vitamins, scarves, clothes, magazines, pizza, salad, bread, wolves, frogs, planes |
| 4 | sent | low | The burglar stole Peter's briefcase, camera, bike, computer, and wallet. |
| 4 | list | low | ginger, figures, hare, blender, briefcase, camera, bike, ham, cows, knot |
| 4 | sent | high | Jeff reluctantly sold his motorcycle, skateboard, bike, van, and cart. |
| 4 | list | high | needles, phone, razors, wrench, motorcycle, skateboard, bike, ribs, silk, mattress |
| 5 | sent | low | The puppy chewed the toy, book, pillow, phone, and ball. |
| 5 | list | low | bulbs, skunk, apron, dentist, toy, book, pillow, passport, plates, hens |
| 5 | sent | high | The maid cleaned the window, blanket, pillow, curtain, and mat. |
| 5 | list | high | rocks, bed, oceans, umbrella, window, blanket, pillow, diapers, soup, stool |
| 6 | sent | low | The handyman repaired his toilet, fan, stove, fridge, and monitor. |
| 6 | list | low | pencil, spoons, corn, sand, toilet, fan, stove, lamps, snakes, gazelles |
| 6 | sent | high | The fire damaged the skillet, oven, stove, chimney, and fridge. |
| 6 | list | high | ketchup, mops, chips, casket, skillet, oven, stove, sage, markets, street |
| 7 | sent | low | The cat knocked over photos, plants, pans, ashtrays, and mugs. |
| 7 | list | low | toaster, sticks, bracelets, pillow, photos, plants, pans, glitter, tobacco, bones |
| 7 | sent | high | Katie clumsily dropped the trays, dishes, pans, tins, and cans. |
| 7 | list | high | ashtrays, cucumber, otters, wraps, trays, dishes, pans, chimney, sponges, fridge |
| 8 | sent | low | The art project requires paint, thread, paper, denim, and glitter. |
| 8 | list | low | beans, carriages, book, arrows, paint, thread, paper, alarm, diaries, parrot |
| 8 | sent | high | Lee put away the glue, ink, paper, tape, and wax. |
| 8 | list | high | chili, husks, bathtubs, disks, glue, ink, paper, robes, corpse, chicken |
| 9 | sent | low | The girl collected several sticks, acorns, shells, bark, and leaves. |
| 9 | list | low | floors, avocado, bows, tin, sticks, acorns, shells, rubber, attic, acorns |
| 9 | sent | high | The collector selected some rocks, pebbles, shells, feathers, and clovers. |
| 9 | list | high | portrait, bishop, nest, carrots, rocks, pebbles, shells, frames, skeleton, camels |
| 10 | sent | low | The eccentric collector hoarded lamps, dolls, guitars, watches, and shoes. |
| 10 | list | low | forks, pen, toilet, rodeo, lamps, dolls, guitars, wood, symbols, straps |
| 10 | sent | high | The music store sells pianos, violins, guitars, drums, and clarinets. |
| 10 | list | high | theater, graves, drums, mulch, pianos, violins, guitars, crates, knuckles, cocoas |
| 11 | sent | low | Hayley carved figurines of pandas, leopards, dolphins, koalas, and bears. |
| 11 | list | low | trunks, rails, cathedral, crickets, pandas, leopards, dolphins, balloons, scarfs, pony |
| 11 | sent | high | Lisa loves talking about whales, sharks, dolphins, otters, and beavers. |
| 11 | list | high | dogs, lizards, shields, broth, whales, sharks, dolphins, parks, bridges, dove |
| 12 | sent | low | The maid prepared some buns, cupcakes, potatoes, prawns, and burgers. |
| 12 | list | low | mugs, batter, ants, mailbox, buns, cupcakes, potatoes, husk, lobsters, gym |
| 12 | sent | high | Paul the farmer grows tomatoes, beans, potatoes, peas, and carrots. |
| 12 | list | high | receipts, ducks, canals, flour, tomatoes, beans, potatoes, puddings, ears, bourbon |
| 13 | sent | low | The stylists decorated every flyer, backdrop, hat, costume, and banner. |
| 13 | list | low | mint, plazas, medals, silver, flyer, backdrop, hat, tins, tables, eagle |
| 13 | sent | high | Ronald had forgotten his bag, suit, hat, tie, and umbrella. |
| 13 | list | high | deer, tortoises, postcards, barks, bag, suit, hat, pigs, peppers, cloak |
| 14 | sent | low | Rob made sketches of farms, oceans, mountains, lakes, and coasts. |
| 14 | list | low | shirt, clocks, almonds, moose, farms, oceans, mountains, colt, jackets, spinach |
| 14 | sent | high | Adam likes hiking through hills, deserts, mountains, valleys, and rivers. |
| 14 | list | high | wrappers, pants, infant, lavender, hills, deserts, mountains, chocolate, watch, bars |
| 15 | sent | low | The mad scientist studied crickets, goats, birds, donkeys, and toads. |
| 15 | list | low | lids, pouches, horses, loaf, crickets, goats, birds, airport, mules, stumps |
| 15 | sent | high | The scientist adopted various rabbits, snakes, birds, cats, and lizards. |
| 15 | list | high | elk, creeks, backpack, wax, rabbits, snakes, birds, garage, hallways, soda |
| 16 | sent | low | The comedic story featured frogs, horses, mice, hens, and ducks. |
| 16 | list | low | donkeys, mustard, zebras, foreheads, frogs, horses, mice, strawberry, olives, centers |
| 16 | sent | high | The infested shed has rats, bugs, mice, roaches, and worms. |
| 16 | list | high | cake, coffee, cats, peaches, rats, bugs, mice, brushes, spaghetti, helmets |
| 17 | sent | low | The ecologist wrote about lizards, camels, foxes, gazelles, and oases. |
| 17 | list | low | canes, dinosaurs, clovers, badges, lizards, camels, foxes, molds, scrolls, tutor |
| 17 | sent | high | While trekking Clara spotted squirrels, wolves, foxes, owls, and deer. |
| 17 | list | high | ox, freeways, lace, toothbrush, squirrels, wolves, foxes, rosemary, subway, elbow |
| 18 | sent | low | The child read about pigeons, turtles, giraffes, penguins, and seals. |
| 18 | list | low | bench, burgers, sleeves, oil, pigeons, turtles, giraffes, soap, jasmine, sofas |
| 18 | sent | high | The zookeeper fed the elephants, monkeys, giraffes, lions, and gorillas. |
| 18 | list | high | axes, shovel, toes, combs, elephants, monkeys, giraffes, butterfly, socks, plug |
| 19 | sent | low | The children had some juice, kiwi, cereal, cinnamon, and grapes. |
| 19 | list | low | stamp, garden, liver, phone, juice, kiwi, cereal, candles, rivers, coasts |
| 19 | sent | high | For snacks Roman prefers vegetable, fruit, cereal, chocolate, and berries. |
| 19 | list | high | button, jewels, lions, marbles, vegetable, fruit, cereal, thread, pies, cashier |
| 20 | sent | low | The dressing recipe requires honey, oil, pepper, vinegar, and mustard. |
| 20 | list | low | swamps, clay, monitor, alligator, honey, oil, pepper, gloves, limes, garlic |
| 20 | sent | high | The cook combined some flour, chicken, pepper, paprika, and ginger. |
| 20 | list | high | gorillas, spices, bottles, elephants, flour, chicken, pepper, butter, ball, jelly |
| 21 | sent | low | Dan is growing some lavender, sage, parsley, jasmine, and oregano. |
| 21 | list | low | pancakes, wine, trays, toads, lavender, sage, parsley, popcorn, wheats, graphs |
| 21 | sent | high | Pete chopped up some thyme, chives, parsley, mint, and basil. |
| 21 | list | high | oven, glue, trains, clarinet, thyme, chives, parsley, entrances, monuments, paint |
| 22 | sent | low | Anna kindly donated her heels, bags, sweaters, belts, and blouses. |
| 22 | list | low | heart, gravy, scorpions, luggage, heels, bags, sweaters, statues, suit, wheelchairs |
| 22 | sent | high | Pat carelessly shrunk her pants, jeans, sweaters, socks, and jackets. |
| 22 | list | high | peacocks, shoes, koalas, spices, pants, jeans, sweaters, figs, stairs, valleys |
| 23 | sent | low | The ancient chest contained skulls, gems, swords, pendants, and pouches. |
| 23 | list | low | doughnuts, pants, flyer, backdrop, skulls, gems, swords, knees, dishes, carpets |
| 23 | sent | high | The mighty warriors yielded spears, bows, swords, maces, and axes. |
| 23 | list | high | pianos, violins, canvas, toilets, spears, bows, swords, brain, buses, pyramids |
| 24 | sent | low | The council built several monuments, plazas, malls, squares, and memorials. |
| 24 | list | low | vendors, cellar, lamb, cranes, monuments, plazas, malls, saffron, penguins, casino |
| 24 | sent | high | The tour stops at stores, restaurants, malls, parks, and shops. |
| 24 | list | high | sculptures, leopards, lipsticks, farms, stores, restaurants, malls, ivory, worms, spines |
| 25 | sent | low | The janitor cleaned the offices, galleries, stairs, entrances, and courtyards. |
| 25 | list | low | lights, shotgun, camera, crackers, offices, galleries, stairs, maid, plants, owls |
| 25 | sent | high | Naughty Andy scribbled on curtains, doors, stairs, windows, and closets. |
| 25 | list | high | theater, shovels, cans, martini, curtains, doors, stairs, compass, cribs, picnic |
| 26 | sent | low | The old woman collects husk, straw, gravel, grass, and hay. |
| 26 | list | low | motorcycle, skateboard, dolls, wool, husk, straw, gravel, bag, tomatoes, kiwi |
| 26 | sent | high | The lone worker carried sand, cement, gravel, wood, and bricks. |
| 26 | list | high | cloth, squirrels, buns, cupcakes, sand, cement, gravel, porcelain, charts, station |
| 27 | sent | low | The antique shop sells mirrors, clocks, goblets, candles, and frames. |
| 27 | list | low | stores, chamomile, ceilings, pizza, mirrors, clocks, goblets, hose, pin, photograph |
| 27 | sent | high | The dragon guarded the vases, jewels, goblets, lockets, and scrolls. |
| 27 | list | high | bananas, costume, peach, nun, vases, jewels, goblets, cheese, claw, vases |
| 28 | sent | low | Merchants used to trade porcelain, china, cotton, saffron, and ivory. |
| 28 | list | low | spiders, flower, bags, restaurants, porcelain, china, cotton, pottery, porch, jars |
| 28 | sent | high | The outfit consisted of wool, linen, cotton, silk, and polyester. |
| 28 | list | high | fruit, skillet, squire, grapes, wool, linen, cotton, runners, closets, rabbits |
| 29 | sent | low | The hiker packed some clothes, matches, maps, cookies, and water. |
| 29 | list | low | thyme, spears, cemetery, camels, clothes, matches, maps, shacks, paprika, couch |
| 29 | sent | high | The numbered plots included graphs, charts, maps, drawings, and tables. |
| 29 | list | high | globe, vest, pitcher, ship, graphs, charts, maps, buildings, memorials, tuxedo |
| 30 | sent | low | Cancer spread to Christine's brain, spleen, stomach, bones, and larynx. |
| 30 | list | low | jaws, thorn, offices, infants, brain, spleen, stomach, mat, dresser, maces |
| 30 | sent | high | The explosion injured Molly's liver, pancreas, stomach, heart, and lungs. |
| 30 | list | high | ribbons, juice, squares, bark, liver, pancreas, stomach, swans, churches, yogurt |
| 31 | sent | low | Artists creatively painted using sponges, brushes, fingers, crayons, and pencils. |
| 31 | list | low | trophy, caps, hangers, paper, sponges, brushes, fingers, china, foam, necklaces |
| 31 | sent | high | Tom accidentally hurt his knees, knuckles, fingers, toes, and ears. |
| 31 | list | high | curtain, straw, notebooks, matches, knees, knuckles, fingers, pasta, biscuits, falcon |
| 32 | sent | low | Tourists enjoyed visiting the markets, stalls, trucks, vendors, and landmarks. |
| 32 | list | low | grains, vinegar, bowls, monkeys, markets, stalls, trucks, tape, rats, mustache |
| 32 | sent | high | The engineers examined the cars, buses, trucks, trains, and ships. |
| 32 | list | high | trophies, insect, wallet, bats, cars, buses, trucks, crayons, fudge, valves |
| 33 | sent | low | Beach visitors left behind wrappers, magazines, bottles, wipes, and containers. |
| 33 | list | low | leash, cafes, boards, cinema, wrappers, magazines, bottles, telescope, balloon, batteries |
| 33 | sent | high | It's important to recycle jars, cans, bottles, cardboard, and paper. |
| 33 | list | high | yarn, window, airplane, cacti, jars, cans, bottles, briefcase, clerk, gems |
| 34 | sent | low | The billionaire acquired every cinema, theater, hotel, casino, and resort. |
| 34 | list | low | petals, pandas, violin, spleen, cinema, theater, hotel, pencils, pendants, card |
| 34 | sent | high | Special forces searched every street, station, hotel, airport, and hall. |
| 34 | list | high | logs, prawns, crab, gutters, street, station, hotel, straws, van, pond |
| 35 | sent | low | The earthquake destroyed many pipes, lights, roads, towers, and reservoirs. |
| 35 | list | low | salad, sandwiches, fountain, shelf, pipes, lights, roads, dolls., berries, meat |
| 35 | sent | high | Urban planners designed the highways, bridges, roads, buildings, and canals. |
| 35 | list | high | drawers, whales, linen, hay, highways, bridges, roads, nuns, chef, peas |
| 36 | sent | low | The famous architect included ponds, fountains, lawns, garages, and pools. |
| 36 | list | low | kangaroos, drawings, dime, torpedo, ponds, fountains, lawns, bartenders, portraits, squids |
| 36 | sent | high | Big mansions often have gardens, porches, lawns, yards, and cellars. |
| 36 | list | high | pancreas, pigeons, cement, tea, gardens, porches, lawns, reservoirs, sheet, bakery |
| 37 | sent | low | The auction featured some desks, carpets, paintings, chandeliers, and frames. |
| 37 | list | low | roaches, heels, rafts, cinnamon, desks, carpets, paintings, rug, florist, sharks |
| 37 | sent | high | The new gallery boasts statues, sculptures, paintings, pictures, and pottery. |
| 37 | list | high | museum, lungs, courtyards, lockets, statues, sculptures, paintings, cats, knife, drums |
| 38 | sent | low | The shopper purchased some olives, figs, oranges, prunes, and squids. |
| 38 | list | low | highways, folder, toy, galleries, olives, figs, oranges, tavern, dill, whisky |
| 38 | sent | high | The fruit basket contains apples, peaches, oranges, bananas, and pears. |
| 38 | list | high | tomatoes, lamp, scalp, bricks, apples, peaches, oranges, oregano, pictures, sidewalks |
| 39 | sent | low | The traveler ate some tuna, avocado, rice, cucumber, and seaweed. |
| 39 | list | low | bugs, photos, plant, ceiling, tuna, avocado, rice, water, ships, leaves |
| 39 | sent | high | Canteen food usually includes corn, fruit, rice, meat, and vegetable. |
| 39 | list | high | raisins, flask, freezer, oases, corn, fruit, rice, cook, belts, skulls |
| 40 | sent | low | The charity gave out vitamins, scarves, shirts, thermals, and shampoo. |
| 40 | list | low | blanket, chests, resort, wrist, vitamins, scarves, shirts, computer, fern, tissues |
| 40 | sent | high | Cleaners bleached the white socks, pants, shirts, towels, and sheets. |
| 40 | list | high | fireworks, coyote, cauldron, fists, socks, pants, shirts, jeans, pipes, cardboard |
| 41 | sent | low | The international company shipped tin, rubber, tea, spices, and sugar. |
| 41 | list | low | grass, lakes, chandeliers, blouses, tin, rubber, tea, hall, seaweed, bears |
| 41 | sent | high | The small cafe sells cake, coffee, tea, sandwiches, and wraps. |
| 41 | list | high | landmarks, denim, shampoo, towels, cake, coffee, tea, blouses, pears, bucket |
| 42 | sent | low | The woodworker specializes in rails, cabinets, doors, frames, and chests. |
| 42 | list | low | larynx, tie, thermals, pill, rails, cabinets, doors, banner, clarinets, rivers |
| 42 | sent | high | The contractor redid the floors, windows, doors, stairs, and ceiling. |
| 42 | list | high | curtains, towers, sheets, banner, floors, windows, doors, prunes, beavers, leaves |

